# Microbiome Maps: Hilbert Curve Visualizations of Metagenomic Profiles

**DOI:** 10.1101/2021.03.22.436520

**Authors:** Camilo Valdes, Vitalii Stebliankin, Daniel Ruiz-Perez, Ji In Park, Hajeong Lee, Giri Narasimhan

## Abstract

**Motivation:** Abundance profiles from metagenomic sequencing data synthesize information from billions of sequenced reads coming from thousands of microbial genomes. Analyzing and understanding these profiles can be a challenge since the data they represent are complex. Particularly challenging is their visualization, as existing techniques are inadequate when the taxa number is in the thousands. We present a technique, and accompanying software, for the visualization of metagenomic abundance profiles using a space-filling curve that transforms a profile into an interactive 2D image.

**Results:** We created Jasper, an easy to use tool for the visualization and exploration of metagenomic profiles from DNA sequencing data. It orders taxa using a space-filling Hilbert curve, and creates a “Microbiome Map”, where each position in the image represents the abundance of a single taxon from a reference collection. Jasper can order taxa in multiple ways, and the resulting *microbiome maps* can highlight “hot spots” of microbes that are dominant in taxonomic clades or biological conditions.

We use Jasper to visualize samples from a variety of microbiome studies, and discuss ways in which *microbiome maps* can be an invaluable tool to visualize spatial, temporal, disease, and differential profiles. Our approach can create detailed *microbiome maps* involving hundreds of thousands of microbial reference genomes with the potential to unravel latent relationships (taxonomic, spatio-temporal, functional, and other) that could remain hidden using traditional visualization techniques. The maps can also be converted into animated movies that bring to life the dynamicity of microbiomes.

**Availability:** Jasper is freely available at microbiomemaps.org and via biorg.cs.fiu.edu/jasper

**Supplementary information:** Supplementary materials are available at microbiomemaps.org

## 1 Introduction and Background

Microbiome samples are routinely processed by means of low-cost, high-throughput metagenomics DNA sequencing, followed by the creation of microbial community abundance profiles [16], where the sequenced reads are mapped against a collection of microbial reference genomes like Ensembl [1] or RefSeq [34]. Tools such as Flint [41] and Kraken 2 [45] facilitate the creation of microbial abundance profiles either from metagenomic wholegenome DNA sequencing (mWGS) or 16S-amplicon sequencing (16S) data. The abundance profiles are the stepping stones for downstream analyses such as differential abundance studies [43], co-occurrence pattern discovery [23, 25, 42, 24], Bayesian analyses [22, 39], biomarker identification [40], multi-omics analyses [8, 9], and analyses of profiles from longitudinal studies [31, 38]. Lower sequencing costs have resulted in an increasing number of larger deep sequencing metagenomic data sets [32].

Metagenomic profiles contain relative abundance values for the entire collection of microbial taxa present in a sample, and these profiles can be easily viewed with stacked bar charts or pie charts using data analysis software suites such as Tableau [7] or MS Excel [5]. Such tools are readily available to the public and allow for data exploration but are designed for the analysis of generic tabular data, and do not consider domain-specific information (taxonomic, phylogenetic, etc.) that may be crucial for the interpretation of metagenomics datasets. Recent tools such as WHAM! [21] allow for explicit metagenomics-focused analyses, making it possible to dig down into the data and create useful visualizations for descriptive analyses.

We argue that complex latent properties of microbiomes embedded in community abundance profiles such as taxonomic hierarchies and other relationships are not easily described and visualized in traditional generic plotting mechanisms such as stacked bar-charts or line-plots. The problem gets more acute as the sizes of our reference genome collections continues to grow exponentially over time [33], and even novel tools such as Krona [35] are not able to keep up and display large amounts of reference genomes.

In this work we consider the problem of visualizing a microbiome using a visualization technique called the *Hilbert Curve Visualization* (HCV). We visualize abundance profiles using the Jasper tool, a free and easy to use graphical application, and discuss the challenges of visualizing billions of measurements for hundreds of thousands of microbial genomes. We propose an alternative visualization technique that is useful when trying to combine many factors of metagenomic information in order to create interpretable images that can lead to improved understanding.

## 2 Approach

We use a technique called the Hilbert curve visualization (HCV) to visualize the microbial community abundance profiles of a reference collection of genomes as a “microbiome map”. For our experiments, we used a reference collection of 44K genomes [1], but the approach is readily scalable to deal with considerably larger collections. These profiles contain the relative abundance measurements of thousands of genomes, and they are ordered along a space-filling curve in a 2D square using the Hilbert curve [28]. Thus, in its simplest form, it is possible to visualize the profile of a single metagenomic sample. In the resulting 2D Hilbert image, each position (or pixel) corresponds to a genome from the reference collection and the position’s intensity color value represents the relative abundance of a single genome in the sample.

As discussed below, depending on the ordering of the genomes that is selected in the software, different “Microbial Neighborhoods” are created, allowing for different interpretations of the “hotspots” of abundant genomes in the images. As explained later, the ordering in Figure 2 allows us to infer taxonomic clades that are most abundant in a sample, while the ordering in Figure 3 allows us to visualize site-specific or stage-specific taxa abundant in a sample.

**Fig. 1.**
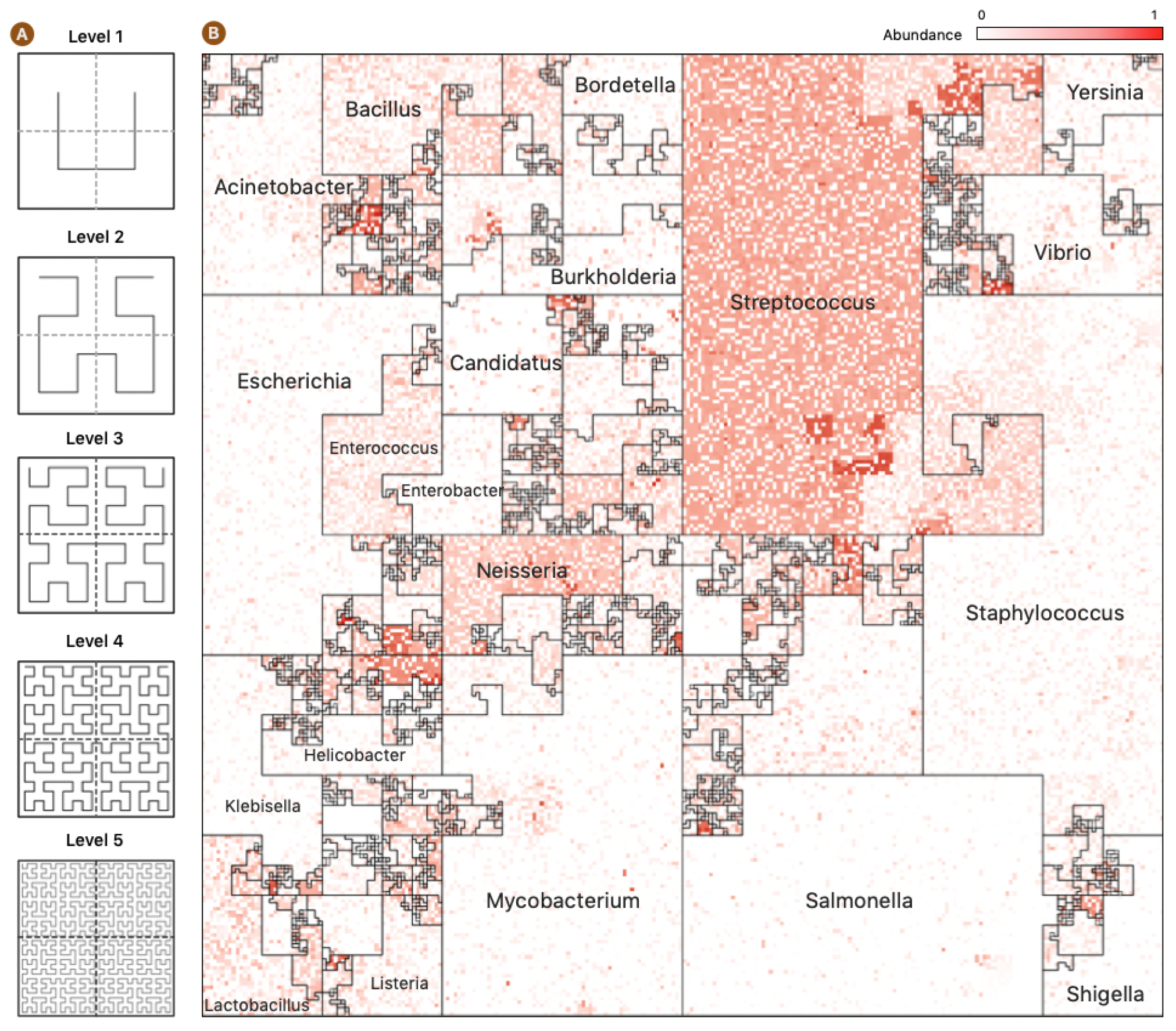
Hilbert Curve Visualization of Metagenomic Samples. **(A)** The first five iterations of the Hilbert curve: the Level 1 curve is obtained by connecting the centers of the four initial squares as shown; the Level *k* curve is obtained by a recursive partitioning of each square from Level *k* – 1, creating four Level *k* – 1 curves and connecting them as outlined by the Level 1 curve, rotated appropriately. At level *k*, the original square is divided into 2^*k*^ × 2^*k*^ small squares, each of whose centers is visited by the Level *k* Hilbert curve. **(B)** A representative image of a mWGS Buccal Mucosa sample from the Human Microbiome Project (HMP) created using a “taxonomic ordering” of 44K reference genomes from the Ensembl database. The color intensity of each position in the image represents the abundance of one microbial genome. The bordered regions are “Microbial Neighborhoods” and represent groups of related microbes, and their size corresponds to the number of genomes in the reference collection.

**Fig. 2.**
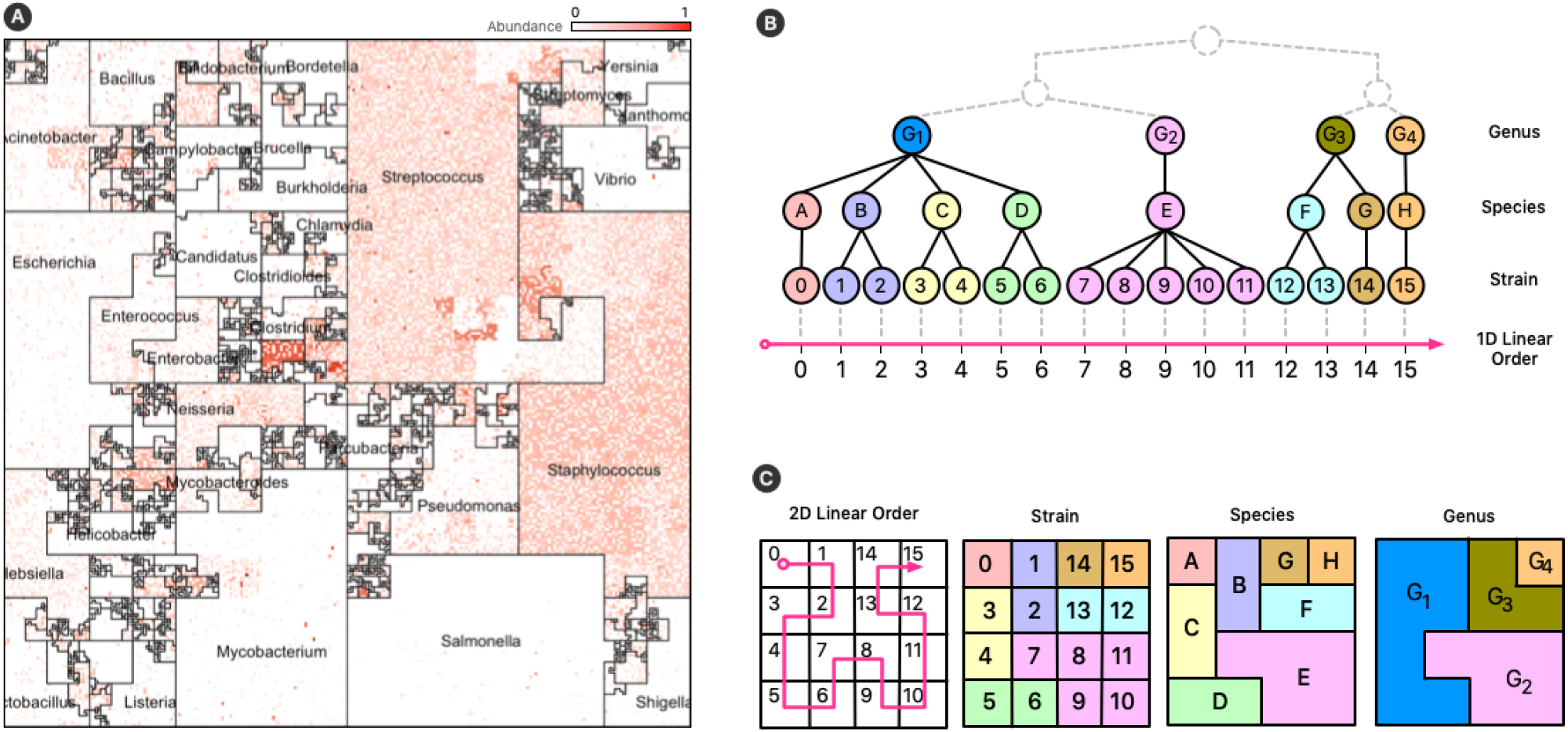
Taxonomic Ordering. **(A)** A *Taxonomic Ordering* of 44,048 reference genomes from Ensembl Bacteria. *Microbial neighborhoods* are drawn based on a taxonomic tree for microbial classification. The image depicts the distribution of Genera in the reference collection, and the size of each neighborhood is consistent to the number of genomes that belong to it. **(B)** Taxonomic hierarchical tree with three levels of rankings for genomes: Genus, Species, and Strain. The tree is linearized to create a 1D linear order for the tree’s leaves (Strains). **(C)** The 1D linear order is laid out onto a 2D plane using a Hilbert curve, creating *microbial neighborhoods* of related taxa.

**Fig. 3.**
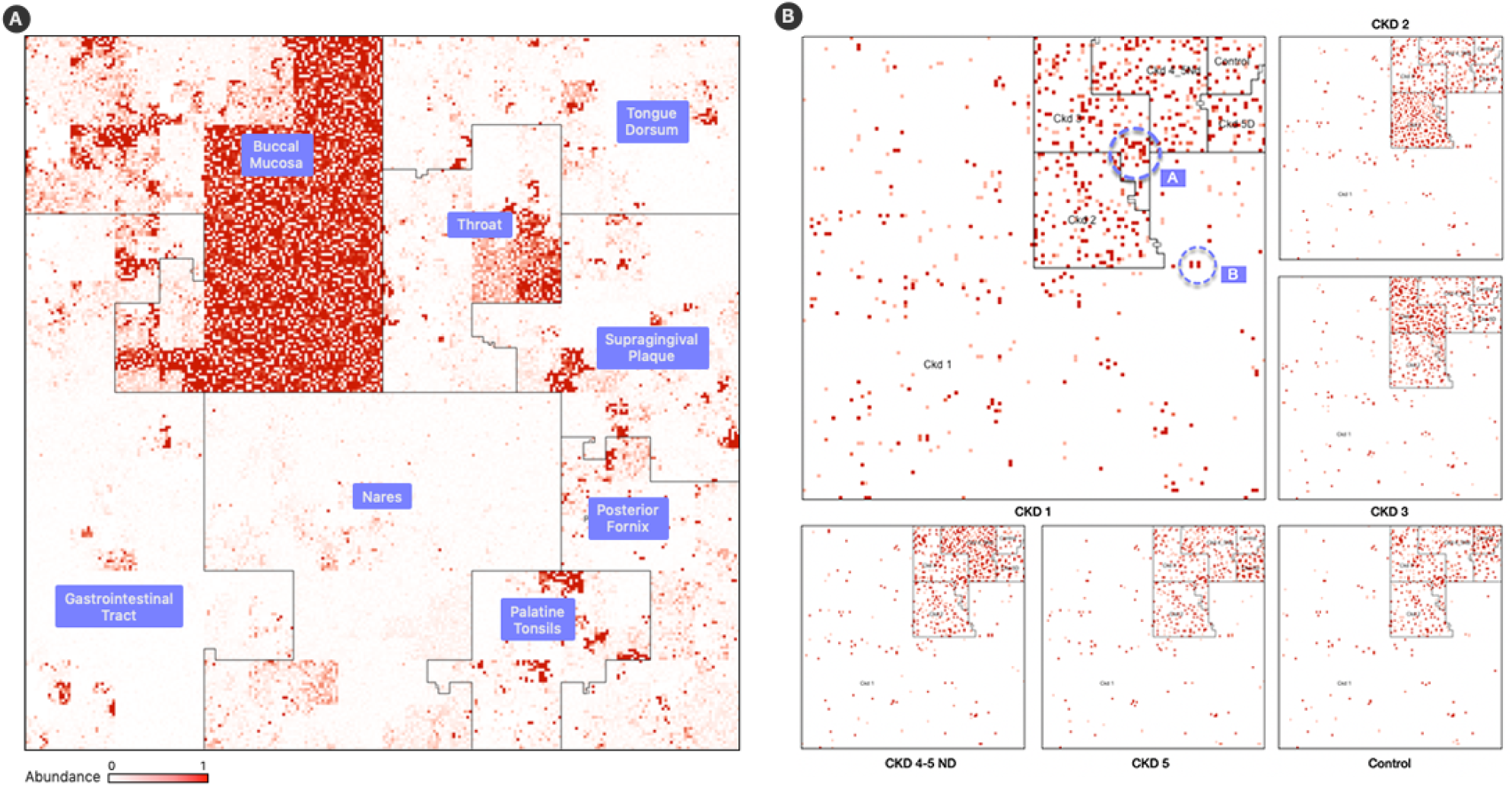
Labeled Orderings. Samples are processed in a *M* × *N* matrix that contains *M* labeled samples, and *N* taxons. In these examples, the taxa with the highest mean relative abundance is identified and used as an anchor for a linear order. **(A)** A map of a buccal mucosa condition from HMP. **(B)** Six maps of representative fecal samples from each CKD stage. The “A” and “B” regions highlighting species which discern the stages.

### 2.1 Space-Filling Curves

Space-filling curves are popular in scientific computing applications for their ability to speed-up computations, optimize complex data structures, and simplify algorithms [14]. Trees are particularly interesting structures that can be optimized with space-filling curves because it is possible to generate sequential orderings of the nodes of the tree in which parent and children nodes are neighbors in a 2D plane. The combination of trees and space-filling curves has been shown to be useful in many fields [12], and in metagenomics this can be useful because the microbial genomes in a reference database are classified using a taxonomy tree with a hierarchy of levels (Strain, Species, Genus, etc.). For data from mWGS experiments, we can presume the leaf nodes of the taxonomy tree to be microbial strains (Figure 2, panel (B)); for 16S data, the leaf nodes are usually species or genera. Clades of the taxonomy tree correspond to microbial neighborhoods in the resulting visualization.

### 2.2 The Hilbert Curve

The Hilbert curve is one of the more prominent examples of space-filling curves, and its construction is based on a recursive partitioning of a square into four subsquares, and then connecting the centers of these squares in a specific order. To provide a recursive definition of the curve, Figure 1 (A) shows the curve at Level 1 when there are only four squares to connect. The curve at Level *k* is defined recursively by dividing the original square into four squares, each with a Level *k* – 1 curve in it and then connecting these pieces using the template of the Level 1 curve after appropriate rotation of the four curves.

Many applications exploit the order that space-filling curves impose on data, and a particular application has been the visualization of high-dimensional data. The first use of the Hilbert curve as a visualization tool was proposed by Keim in 1996 [30] to represent stock market data. Since then, it has been used for visualizing genomic data [19, 10] and DNA alignments of whole bacterial genomes [44].

In human genomics, the application of the HCV technique is straightforward as the natural linear order of genomic positions can be easily used by the curve, and there are tools for creating Hilbert curve images from genomics datasets (HilbertVis [10] and HilbertCurve [27]). Both of these tools apply HCV in the context of human genomics: a single scaffold is modeled as a single one-dimensional (1D) line in which each interval is taken to be a single genomic position. To date, the HCV technique has not been applied to metagenomics datasets.

#### 2.2.1 Visualizing Metagenomics Data

Traditional 1D visualization techniques that display community abundance profiles often do not take into account latent metagenomic factors present in sequencing samples. Pie charts, line plots, etc., tend to focus on the taxa with the highest abundances, and have poor resolution for taxa with small abundances (which sometimes can be critical). Visual comparison of multiple samples is also difficult as determining the change in abundance of a single taxon between samples is not convenient.

Space-filling curves offer an intriguing scheme for visualizing metagenomics data for their ability to preserve positional data. This feature can be enhanced with a reference collection’s metadata to create descriptive images that express metagenomic information succinctly. The issue of adding new genomes is discussed in detail in Section 3.2.

The Hilbert curve is not the only space-filling curve with these features, but its creation is a simple recursive partitioning that can be implemented elegantly and efficiently in software.

Other curves, like the Peano curve [36], partition the square into 9 or more regions, which can lead to hard to interpret images when the levels of the curve are high (levels of 10 or above).

#### 2.2.2 Linear Orderings

The first challenge in visualizing abundance profiles with a space-filling curve is that there is no natural linear order for the reference collection. Below, we discuss two classes of orderings that are shown to be useful:

- **Taxonomic Ordering:** Based on a taxonomic tree.
- **Labeled Ordering:** Based on a custom labeling scheme.

The area of the 2D square that bounds a *microbiome map* is proportional to the number of unique genomes in the reference collection. For a curve of size *k*, we have 2^*k*^ × 2^*k*^ linear segments in which to place a genome in. However, the size of a reference collection will not always perfectly match the number of segments. To account for this, we merge adjacent segments along the curve’s path so that their number is always equal to the number of genomes in the reference collection. Jasper has controls to understand a map’s linear order: users can overlay the path of the curve to follow the map’s layout (Figure 4).

**Fig. 4.**
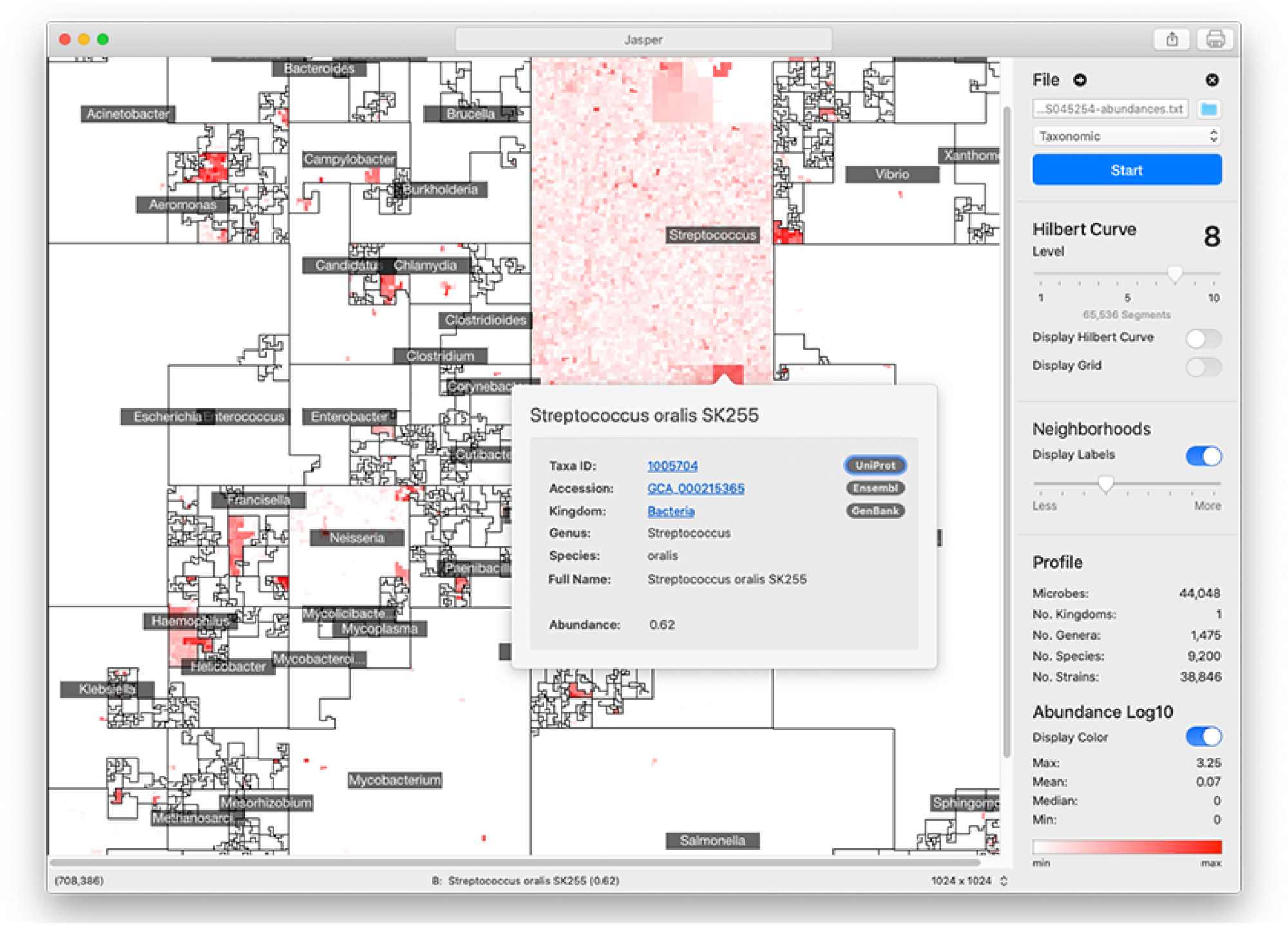
Jasper User Interface. The Jasper software is a tool for interactively exploring *microbiome maps*. Users can click on any genome in the map and get a pop-over with detailed information, along with links to specific online resources about the genome in Ensembl, Uniprot, and Genbank. The maps can be exported and easily shared.

## 3 Methods

Visualizing a microbiome’s abundance profile starts by aligning DNA sequencing reads against a reference collection of genomes, and creating counts of the number of reads that align to each genome. The cloud-based tool called Flint [41] facilitates the profiling of mWGS datasets, while the Kraken 2 software does it for 16S datasets. For the images shown in this paper, Flint uses a reference collection of 44,408 microbial genomes from the Ensembl Bacteria database [1], while Kraken 2 uses a “16S” reference of 5,127 genomes which contains references from Greengenes [20], SILVA [37], and RDP [17].

Different linear orderings of the taxa on the Hilbert curve result in different images. In Jasper, users may select from two options: a *taxonomic ordering* (Figure 2) which uses the linear order from Ensembl’s taxonomic tree, or a *labeled ordering* (Figure 3) which is based on a custom user-supplied label. The project website at microbiomemaps.org contains a manual with examples that show how to create them. Note that unlike other HCV techniques, *microbiome maps* do not depict genomic positions: each pixel region corresponds to the abundance of a single genome from a reference collection.

### 3.1 Microbial Neighborhoods

Different linear orderings produce different Hilbert curve visualizations, with each resulting in clusters of related microbes along neighboring regions in the 2D plane. The clustering creates unique areas that resemble community neighborhoods in popular consumer mapping applications like Google Maps [3], and we term these areas “*Microbial Neighborhoods*” (Figure 2, panel (A)) as they represent microbes belonging to either the same taxonomic group, or the same biological condition – the idea being that they are clustering around a common scheme. These neighborhoods offer a quick and visual way to readily identify abundance “hotspots” that can contextualize the important features of a metagenomic sample and identify important microbial groups.

#### 3.1.1 Taxonomic Neighborhoods

The first option for ordering genomes along the Hilbert curve is the *taxonomic ordering* which determines a 1D linear order based on a genome’s taxonomic lineage. In this ordering, pairs of taxa belonging to the same taxonomic group (say the same Genus or Species) are placed close to each other along the curve, and consequently, remain generally close to each other in the Hilbert image. This ordering scheme creates “*Taxonomic Neighborhoods*” that envelop related taxa based on their taxonomic lineage, and as seen in Figure 2, multiple taxonomic levels can be displayed at the same time in a single image. The ability of a *microbiome map* to display multiple levels of a taxonomic tree at once, while at the same time providing high-resolution abundance information for single genomes, is a compelling advantage over visualizing data with 1D methods were the sheer number of data points would overwhelm the observer.

An advantage of the *Taxonomic Ordering* is that it creates a visual way of depicting the number of reference genomes in the reference collection, making it possible to compare relative sizes of clades: the *Microbial Neighborhoods* in this ordering represent large clades with many taxa, which in turn occupy large areas of the map. An example of this case is Figure 1 (B) and 2 (A) which shows that the Ensembl bacterial database contains large numbers of strains from the *Streptococcus* and *Staphylococcus* genera.

Another advantage of this ordering is that we can quickly understand the diversity of a reference collection by creating *microbiome maps* with no color; these colorless maps create a visual representation of the reference collection that shows us its taxonomic distribution.

##### Linearizing Taxonomic Trees

Illustrating a taxonomic tree as a 2D Hilbert curve starts by finding a linear order of the leaf nodes in the tree. Figure 2, panel (B), depicts a fictitious taxonomic tree with 16 microbial strains at the leaf nodes ordered along a 1D line using a *taxonomic ordering* scheme (Section 3.1.1) which groups the 16 strains according to their parent species and genus groups. Figure 2, panel (C), illustrates how the 16 strains would be laid out on a 2D plane and how the taxonomic hierarchies are represented as strain, species, and genus areas in the Hilbert image.

Note that tree structures do not have a natural linear order (no “start/finish” or “left/right”. Different tree orderings result by permuting the children nodes at any given node of the tree. Algorithms for finding an optimal order have been proposed [13], but the optimal order relies on the tree satisfying specific properties (e.g., a binary tree) and the existence of a good optimality measure. The *taxonomic ordering* linearizes a tree by using data from Ensembl’s Pan-taxonomic Compara [6] and the Ensembl Genomes [2] databases as the foundation for the Hilbert curve. The genomes in the database are annotated so that we can establish a linear order. For mWGS data, the leaf nodes of the tree are typically at the strain level (for 16S data, the leaf nodes are at the species or genus level). As shown in Figure 2A, the 2D square that bounds the Hilbert image represents all genus-level groups. However, depending on the data and the application this could be modified to suit the needs so that it represents a good compromise between taxonomic information and visual interpretability.

#### 3.1.2 Condition Neighborhoods

The second option for ordering genomes along the Hilbert curve is the *labeled ordering* (Figure 3) which creates “*Condition Neighborhoods*” by using an ordering scheme that determines the 1D linear order based on a user-supplied labeling of samples. This labeling is provided as a labeled *m* × *n* sample matrix *M*, where *m* are sample rows, and *n* are the genomes in the reference database. For sample *i* and reference genome *j*, the matrix entry *M*[*i*, *j*] corresponds to the abundance of genome *j* in sample *i*.

Establishing the linear order for multiple conditions starts with a user-defined ordering of the set of *k* conditions, *C*_1_, *C*_2_,…, *C_k_*. Conditions may represent different disease stages, time intervals, drug dosage, or sampling locations (body sites, environmental sites, etc.).

Once we have a condition ordering established, the next task is to identify taxa whose average relative abundance is highest in *C*_1_ and order them first, followed by taxa whose average relative abundance is highest in *C*_2_, and so on, until we terminate the ordering by taxa that are not abundant in any of the conditions. Once we have established the ordering, we can then draw the Hilbert images for each of the samples from the input sample matrix *M* according to the established order.

Assigning a color to a taxon based on abundance is only meaningful if its presence is above the threshold of noise, which we determine when we normalize the input matrix *M*. In general, if a taxon is most abundant in multiple conditions (something that we have not seen in practice), then we assign it to the first condition as determined by the ordering criteria. After the conditions have been organized along the curve, taxonomic information is used to order genomes within the 2D region of the condition.

In this ordering, the *microbiome map* is still visualizing only one sample, but one can readily spot the relevant biological condition with which it has the most overlap. “*Hotspots*” will most likely appear in the region corresponding to one of the conditions, and users can readily tell what condition the sample belongs to by identifying the area, i.e., neighborhood, in the image with the most hotspots. Thus, it is easy to infer by visual inspection that Figure 3 panel (A) is with high probability a sample from the buccal mucosa region with some taxa that are typically abundant in the throat and other oral sites. Similar inferences are possible with the disease stage or with the environmental condition of the sample. Clusters of bright positions will also appear in other neighborhoods (Figure 3, panel (B)), as other conditions will contain taxa with high relative abundances, but not in the same quantities as for the condition that the sample belongs to.

The process for defining a new custom order is simple. Detailed instructions and examples can be found at the project website at microbiomemaps.org.

### 3.2 Adding New Genomes or Samples

When adding new samples, preserving a genome’s locality or neighborhood becomes important for drawing consistent and useful conclusions about changes. This is easy for the taxonomic ordering. For the labeled ordering, the preservation of locality for a genome is a little more delicate as the assignment of a genome to a neighborhood is done by identifying taxa whose mean relative abundance (or other user-defined metric) is highest in a group of samples. If the new sample to be added changes a taxa’s mean relative abundance in the sample’s group, then it could affect the map’s topology. This is a disadvantage to all mapping orders that rely on precomputing a value to place taxa in a neighborhood – especially when that value is computed across a group of samples, as new samples will require their re-evaluation. Inserting new genomes to the reference collection could affect all existing plots because the positions of existing genomes may change. The addition of new taxa into an existing ordering will also change the relative abundance of all or almost all taxa, changing the color intensities of the pixels. Several suggestions to alleviate the aforementioned problems are provided in Section 4.

## 4 Results and Discussion

We created microbiome maps for two groups of metagenomic datasets: 24 mWGS normal samples taken from the Human Microbiome Project (HMP) [29], and 18 fecal samples (16S) from a collaboration with Kangwon National University and Seoul National University in Korea. The 24 samples from HMP represent 8 different body sites, and the 18 samples from the Korea study represent 5 stages of Chronic Kidney Disease (CKD), along with a normal control set. We analyzed the mWGS HMP samples with the Flint software [41], and the 16S CKD samples with Kraken 2 [45]. For the HMP samples, the metagenomic profiles contained relative abundance measurements for 44,048 microbial strains, and for the CKD samples, the metagenomic profiles contained relative abundance measurements for 5,127 microbial species.

Three samples were selected for our study from HMP from each of the eight following body sites: Buccal Mucosa, Gastro-Intestinal Tract, Nares, Palatine Tonsils, Posterior Fornix, Supragingival Plaque, Throat, and Tongue Dorsum.

Eighteen fecal samples were obtained from CKD patients of Kangwon and Seoul National University Hospitals. The samples were selected based on their glomerular filtration rate (see Kidney Disease Improving Global Outcomes (KDIGO) [4]), and a total of six groups were created: Control, CKD Stage 1 (CKD 1), CKD Stage 2 (CKD 2), CKD Stage 3 (CKD 3), CKD Stage 4 & 5 non-dialysis dependent (CKD 4-5ND), and CKD Stage 5 dialysis dependent (CKD 5). The CKD stages were determined based on the deteriorating function of the patient’s kidneys, and three samples from each group were used.

### 4.1 Microbiome Maps

*Microbiome maps* for mWGS and 16S data communicate abundance information at different levels of a genome’s lineage: for mWGS samples, each position in the image displays information about microbial strains (the resolution at which abundances are reported by Flint [41]); for 16S samples, each position in the image displays information about microbial species (abundances as reported by Kraken 2 [45]). Both sets of profiles were then converted into *microbiome maps* using the *taxonomic*, and *labeled* orders.

Figure 2, panel (A), contains a representative image from one sample of the HMP dataset (Nares) ordered using the *taxonomic ordering* scheme. In this image we can clearly see that the *Streptococcus* and *Staphylococcus* groups are abundant in the Nares sample. While the dominant group would have been obvious even in a traditional 1D plot, the Hilbert curve visualization ensures that the smaller taxonomic groups are not overshadowed by the more abundant groups. Identifying the most abundant taxonomic clades in a sample only takes a quick glance at the image.

Figure 3, panel (A), contains a map from a Buccal Mucosa sample of the HMP dataset. The map contains the same 44K genomes from Figure 2, panel (A), but ordered with a *labeled ordering* scheme, based on the highest mean relative abundance of a genome in its cohort. The advantage of this scheme is that identifying the biological condition that the sample belongs to is effortless: one need only to look at the neighborhood that contains the most *hotspots* (Buccal Mucosa, in this case).

Figure 3, panel (B), shows six maps of 16S samples from the CKD analysis created using the *labeled ordering* scheme, displaying the abundances for 5,127 species each. The order is based on the mean relative abundance of the microbes in the samples belonging to each CKD stage: the most prominent taxa in Stage 1 are surrounded by the CKD 1 area, the most prominent taxa in CKD Stage 2 are surrounded by the CKD 2 area, and so on. Note that the more noticeable regions with a higher density of brighter regions are from samples of the same cohort and we can readily identify the CKD stage by looking at the density of *hotspots*.

What is more significant is the way these plots show the microbes shared by different stages of the disease. For example, the region marked *A* in Figure 3, panel (B), shows a group of microbes that appear in all stages, while the region marked *B* appears in almost all stages except CKD3 (absent) and Control (lowered abundance).

By fixing the orderings of the taxa, a *microbiome map* can be used to present groups of metagenomic samples that can be partitioned temporally (longitudinal studies), spatially (body or environmental sites), by disease (sub)type, by disease stage, and by developmental stages. Additionally, it is readily possible to create *average microbiome maps*, *aggregate* maps, and *differential* maps showing either average, aggregate, or differential abundances, respectively.

To address the problem of adding new genomes or samples, we offer the following suggestions. One possible solution is to leave “blank” (i.e., unassigned) pixels on the map to allow for future additions. This could be implemented by inserting the gap so that the next clade always starts at the boundary of a square region of size 2^*k*^ × 2^*k*^, for a predetermined value of *k*. Second, different parts of the ordering (e.g., a taxonomic clade or a condition associated with a label) can be assigned different colors instead of the monochromatic plots shown here. Finally, it may be useful to always provide a reference microbiome map to clarify the labeling.

#### Comparison to Other Methods

Using 1D visualizations are helpful for condensing information for multiple samples. However, they lack the ability to display nuanced information and to scale to deal with the exponential growth in the databases (44K bacterial strains were used in our visualizations) while retaining the perspective of the latent metagenomic relationships. The project website contains samples visualized with *microbiome maps* and compared to WHAM! and iMAP images.

### 4.2 Jasper Software and Use Cases

Jasper (Figure 4) offers multiple controls for inspecting *microbiome maps*, and integrates with online resources like Ensembl [2], GenBank [15], and UniProt [18]. In Jasper, users can identify any genome in the map by hovering their mouse pointer or clicking on any region. If a user does click, a pop-over is displayed with direct links to the online resources mentioned above, which provide detailed information about the genome. To make the map’s layout easier to understand, the software also allows users to overlay the path of the Hilbert curve on top of the map. Jasper also includes other tools for researchers, as well as the ability to export maps as high quality vector images. The software can also be used with no profiles to create Hilbert curves which can be used for learning about space-filling curves.

Machine learning methods can also benefit from *microbiome maps*, as their unique patterns can surface untapped biological signals that could be used for powerful classification methods.

Jasper is currently localized for North America and English-speaking users. We are working on adapting it to other languages and regions, and also working on accessibility features to make the software easier to use for users with disabilities. Jasper is developed with the Swift programming language [11], and is free to use with no restrictions.

#### Animated Microbiome Movies

*Microbiome maps* can be animated to show the fluctuations of microbe abundances across samples and time. Figure 5 contains a movie strip created from a study by [26] which shows how the microbiome of a single patient changes over time. One can see how the composition is affected by an antibiotic on day 35, and how it recuperates (days 38 - 64). More examples are available at the project website at microbiomemaps.org

**Fig. 5.**
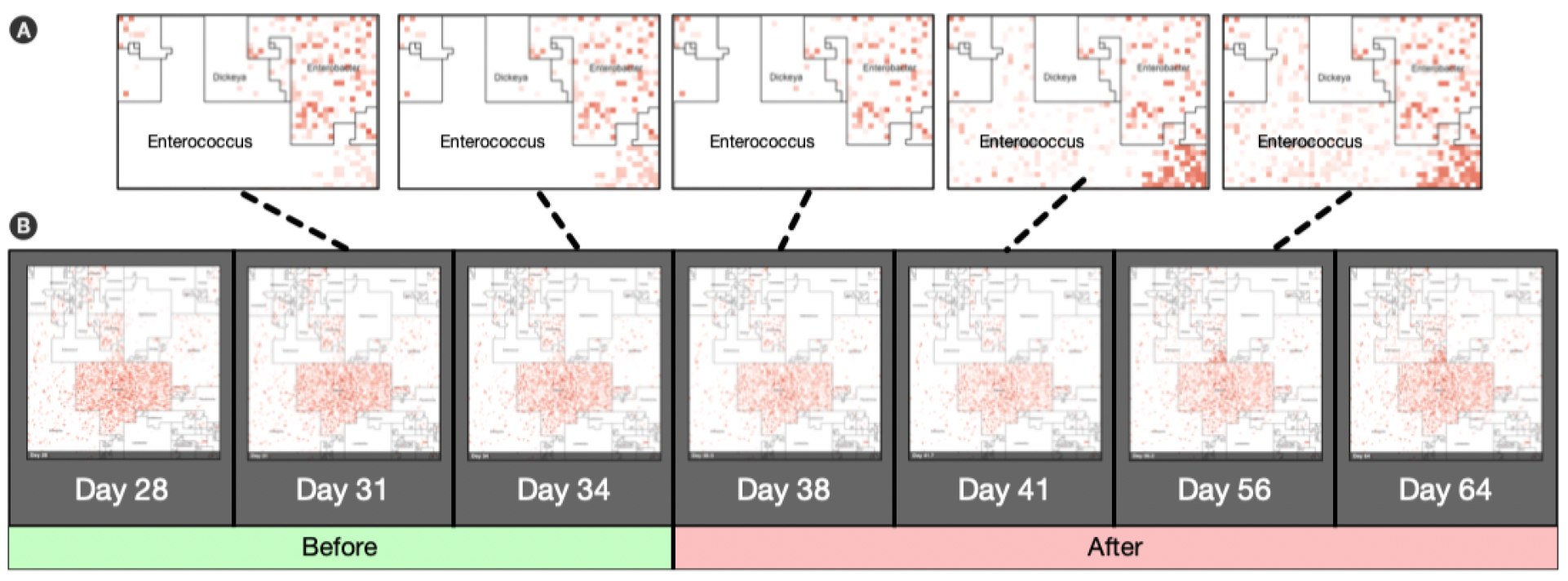
Animated Microbiome Map. Selected frames of an animated time-series visualization of 12,116 strains for a single patient from [26].**Panel (A)**: zoomed regions of the *Enterococcus* neighborhood as it progresses through the antibiotic response. **Panel (B)**: A sample of some animated microbiome maps is available at microbiomemaps.org

#### Project Website: microbiomemaps.org

Jasper is available at microbiomemaps.org. This website is being developed into an online community resource for cataloguing maps. Future releases of Jasper will allow users to share their maps with the website directly from the app, and also on social media. The website will eventually offer curated collections of maps which will be free to use by the community.

## 5 Conclusion

In this work we have shown how the Hilbert curve visualization technique can be used to visualize metagenomic community abundance profiles from both mWGS and 16S DNA sequencing datasets. The resulting *microbiome maps* display the relative abundance of microbial genomes in an interpretable manner, and can convey information about multiple latent factors of the reference genomes in the samples under study.

The Hilbert curve is used to lay out the abundance of microbial taxa from a reference collection using two ordering schemes that can be used to create a *microbiome map*: the first, the *taxonomic ordering* is a default ordering that relies on taxonomic information, and can be used to create images that express abundance values in the context of taxonomic clades that the genomes belong to. The second, the *labeled ordering*, is dependent on a user-specified labeling of biological conditions, and can express the abundance values of the profile in the context of a biological interpretation for a set of samples. Although the above two orders are the first ones to be available in Jasper, we are exploring other orderings that will be incorporated in future releases, such as orderings specific to time-series analyses, or multi-omics datasets.

## Acknowledgments

The authors would like to thank the members of the Bioinformatics Research Group, BioRG, at Florida International University (FIU) for their valuable feedback and comments. We also thank Dr. Jennifer Clarke at UNL, and Dr. Kalai Mathee at FIU, for their helpful comments and feedback on the project.

